# Genetic manipulation of the optical refractive index in living cells

**DOI:** 10.1101/2020.07.09.196436

**Authors:** Junko Ogawa, Yoko Iwata, Nina U Tonnu, Chitra Gopinath, Ling Huang, Sachihiko Itoh, Ryoko Ando, Atsushi Miyawaki, Inder M Verma, Gerald M Pao

## Abstract

The optical refractive index of cellular components is generally not a property considered amenable to manipulation in microscopy as this is an intrinsic physical property of materials. Here we show that by targeting cephalopod reflectin protein nanoparticles one can manipulate the optical refractive index of mammalian cellular compartments. We further demonstrate that refractive index alteration based contrast agents can be utilized for dark field microscopy and quantitative phase contrast holotomography. Additionally we have molecularly cloned novel reflectins with improved and novel optical properties.

## Introduction

Reflectins proteins are the main components responsible for the structural coloration and iridescence in cephalopods (1). This is achieved through specialized cells, the iridophores, which form Bragg mirrors by alternating layers of high and low refractive index material (2). Within the membrane stacks, reflectins form nanoparticles of < 5nm to >500nm and confer the high optical refractive index property to iridophores (3). Reflectins are also utilized by cephalopods in addition for white light scattering and to achieve chromatic invariance of chromatophores, pigment sacs, as these expand or contract (4)(5). The protein consists of multiple reflectin repeats joined by linkers (6). Their amino acid composition is highly unusual and is rich in methionine, arginine as well as aromatic amino acids (1). However they are largely devoid of aliphatic amino acids, lysine and threonine (6)(1). As expected from the lack of aliphatic amino acids, circular dichroism and other global measures of protein structure show them as being highly disordered (7).

## Results

We have explored the idea of using of reflectins as a means to genetically manipulate the refractive index of living cells and specifically their use as novel optical probes based on changes in the local refractive index. To efficiently measure changes in the refractive index in living or fixed cells, we have developed an assay using fluorescence aided cell sorting (FACS). The assay consists of targeting reflectin proteins into the endoplasmic reticulum (ER) of cells. Given that the ER is large and irregular, filling the ER luminal space with a high refractive index material is expected to yield and increase in side scatter. Therefore we synthesized known cephalopod reflectin genes and cloned new reflectin genes sampled from the wild (*Sepioteuthis lessoniana*, Miura peninsula Shiro-ecotype (8)) expected to yield higher refractive index reflectins (Fig. 1A). Reflectin cDNAs synthesized using human codon usage were epitope tagged for antibody recognition with either an HA or FLAG tag. We also inserted a preprotrypsin signal sequence to target the protein into the secretory pathway (Fig. 1A). Addition of a C-terminal KDEL ER retention signal was used for specific ER targeting.

**Figure 1.**
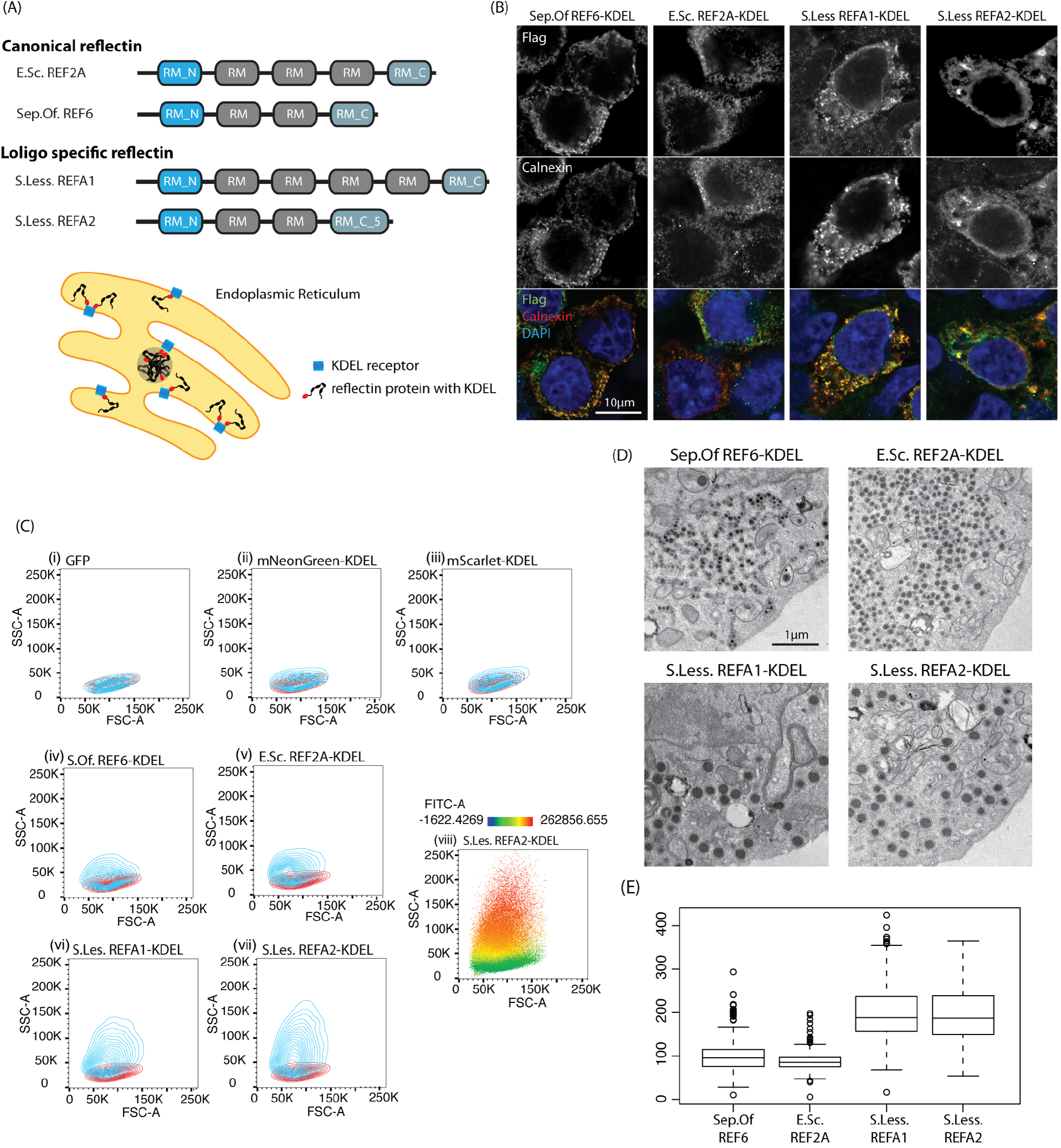
(A) Domain structure of the reflectin proteins used in this study. Reflectins were targeted to the lumen of the ER by addition of a signal sequence and a C-terminal ER retention signal. (B) Immunohistochemical confirmation of ER targeting of reflectin proteins. Reflectin signals colocalize with calnexin an ER marker. (C) A FACS assay for reflectin optical activity: Reflectins increase cellular side scatter (SSC) when targeted to the ER. Scattering is proportional to reflectin expression (viii). *Sepioteuthis lessoniana* Reflectins A1 and A2 have much greater optical activity. (D) Transmission electron microscopy of reflectin nanoparticles in the ER of HEK 293T cells. (E) Size distributions of reflectin nanoparticles produced in human cells. E.sc (*Euprymna scolopes*), Sep Of/S.Of (*Sepia officinalis*), S. Less (*Sepioteuthis lessoniana*)

Immunolocalization of reflectin using the FLAG tag and the ER marker calnexin confirmed colocalization in transiently transfected cells (Fig. 1B).

Reflectin cDNAs in the pBOB-CAG vector were transfected into human embryonic kidney 293 or 293T cells and live cells were evaluated for forward scatter (FSC) vs. side scatter (SSC) by FACS. Control proteins, cytoplasmic GFP and ER targeted mNeonGreen or ER mScarlet did not significantly alter scattering (Fig. 1C ii-iii) when compared to untransfected cells. In contrast, transfection of the cuttlefish (*Sepia officinalis*) reflectin-6 (9) or the Hawaiian bobtail squid (*Euprymna scolopes*) reflectin-2A (6) targeted to the ER yielded a substantial increase in scattering as would be expected if the ER increased in refractive index (Fig. 1C iv-v). The Big Fin Reef squid (*Sepioteuthis lessoniana*) skin has unusually reflective iridocytes, which led us to clone their reflectin genes. Transfection of the *Sepioteuthis lessoniana* ER targeted reflectins A1 and A2 genes yielded substantial increases in scattering, consistent with a possible increase in refractive index within the ER (Fig. 1C vi-vii). As the protein was tagged, we could quantify its abundance in the transfected cells by indirect immunofluorescence. The result shows that expression levels of reflectin A2 correlated with scattering, with the highest reflectin expressing cells displaying the strongest side scatter (Fig. 1C viii). Although scattering is consistent with an increase in refractive index, scattering could also be accomplished by aggregation of large intracellular particles larger than the wavelength of light. Since visible light is in the range of 380-740 nm, we imaged by transmission electron microscopy to ascertain the size of the reflectin nanoparticles. Transmission electron microscopy imaging of ER targeted reflectins showed that they form small spherical osmiumphilic nanoparticles that fill the ER lumen (Fig. 1D). The vast majority of reflectin nanoparticles are well below the wavelength of light with a median diameter of ~100 nm for Euprymna and Sepia reflectins and ~180 nm for *Sepioteuthis* reflectins (Fig. 1E). Only the *S. lessoniana* Reflectin A1 very rarely approached >350 nm in size (1.8%) The *Euprymna scolopes* R2A protein produced the most uniform particle size distributions. These observations are consistent with the view that scattering observed in FACS experiments is not due to simple protein aggregation but due to an increase of the refractive index of the ER lumen.

To test if reflectin targeting could be used as a microscopy label, we tested ER targeted reflectin in dark field microscopy where only reflected or scattered light can be seen.

Fig. 2A shows that all transfected ER targeted reflectins displayed strong darkfield signals, well beyond that of transfected cells of ER targeted mNeonGreen (Fig. 2B). To ascertain that the observed signal was indeed the targeted reflectin, we fused mNeonGreen to either the NH2 or COOH terminus of the *Sepioteuthis* reflectin A2 and transfected substoichiometrically with reflectin A2 (Fig. 2B). In this manner we could ascertain that the mNeonGreen fluorescence and the darkfield signals were completely correlated demonstrating that the darkfield signal indeed originated from reflectin. Quantitative phase contrast microscopy is also sensitive to changes in refractive index, and therefore modifying the local refractive index should produce a robust signal detectable by quantitative phase contrast methods. Moreover if the pathlength is also known, this would allow the direct measurement of the local refractive index. This is indeed what is possible with holotomography, which generates a tomographic reconstruction of the sample. We used holotomography to observe and quantify the change in refractive index within the ER lumen after reflectin targeting (Fig. 2C). In untransfected controls or mScarlet transfected controls, nucleoli are the only observable signal above a refractive index of a RI>1.36. However, signals well above RI of1.36 can be seen in reflectin transfected cells in locations consistent with the ER. High refractive index signals > RI 1.42 could be observed in most cases. The highest refractive indices were achieved with the *Sepioteuthis lessoniana* A2 reflectin and Reflectin 6 from *Sepia officinalis* (data not shown). The holotomography observations were performed in both transfected living cells as well as fixed cells and directly demonstrate the alteration of the optical refractive index within cellular compartments of living mammalian cells.

**Figure 2 Legend.**
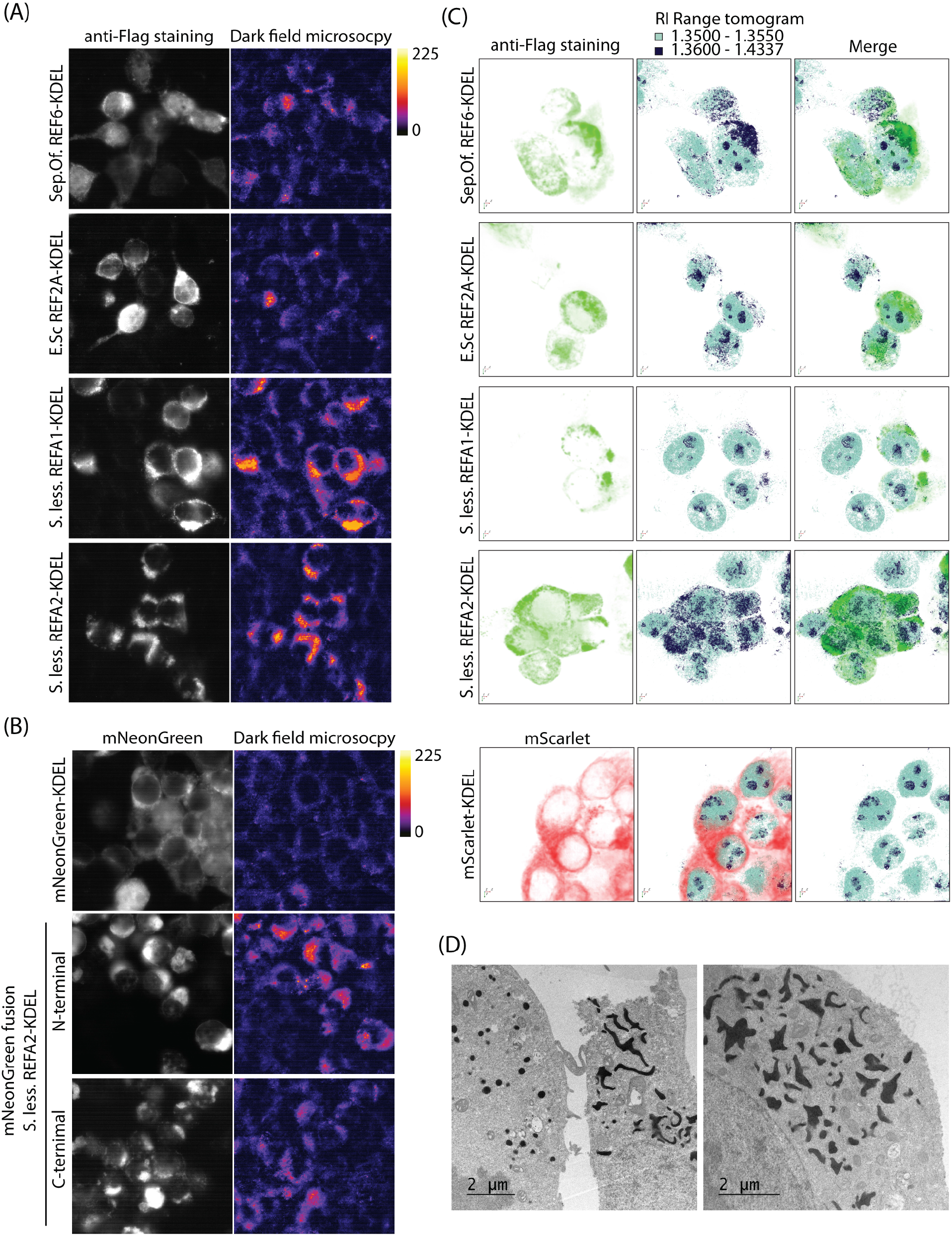
(B) Darkfield microscopic signals of the 4 reflectin proteins targeted to the ER (B) Control mNeonGreen targeted to the ER does not appreciably increase scattering. N- and C-terminal mNeongreen reflectin fusion proteins colocalize with darkfield signals. (C) Direct measurement of refractive index increases in reflectin transfected cells by holotomography. Refractive index increases achieved were over RI >1.4. Localization of mScarlet to the ER did not increase the RI. (D) Mutually exclusive nanoparticle and space filling states of *S. lessoniana* reflectinA2 in the ER lumen. E.sc (*Euprymna scolopes*), Sep Of/S.Of (*Sepia officinalis*), S. Less (*Sepioteuthis lessoniana*)

## Discussion

In summary we report that it is possible to manipulate the optical refractive index of intracellular structures such as the lumen of the ER in living cells using cephalopod reflectins. We demonstrate the utility of this class of proteins as microscopy contrast agents by targeting reflectins into the ER for dark field microscopy as well as quantitative phase contrast microscopy where we observed increases in detectable signal. Using the latter we could demonstrate that we achieved a refractive index of up to ~1.4, well above the RI of cytoplasmic and ER of 1.35. Obviously manipulating the refractive index is probably most useful to achieve transparency of scattering tissues through refractive index matching(10). This requires fixed, permeabilized whole organ samples, which precludes live imaging. However, with a genetically encoded protein this might be conceptually possible.

A large number of challenges, however, remain. To make mammalian tissues transparent, would require bringing intracellular space to approximately a refractive index of RI 1.44 as with the Scale/S technique(11). Given that pure *Euprymna scolopes* reflectin 1A has a refractive index of 1.59 (12), one would require reflectin to encompass 9% of cellular mass, a quantity that might be hard to achieve. In our experiments we achieved a refractive index of ~1.42 albeit locally in small areas and occasional cells in larger areas (data not shown). In addition one would require isotropic distribution throughout the imaged volumes, which is challenging given that reflectins form nanoparticles. This however might be more feasible through reduction of the size of nanoparticles. Ideally one would want an inert liquid of high RI to permeate through intra and extracellular space. Interestingly we have observed that the *Sepioteuthis* reflectin A2 can have two states. It seems to have either liquid-like space filling properties or forms nanoparticles in a given cell but never both in the same cell (Fig. 2D). If refractive index is only a compositional property, the *S. lessoniana* reflectin-A2 or others like it might be the proteins of choice to attempt to achieve in vivo transparency through refractive index matching.

## Materials and Methods

### FACS

Assays were performed on a BD™ LSR II Flow Cytometer and analyses with FlowJo v10.6.

### Immunofluorescence

Immunofluorescence was imaged using a Zeiss LSM 880 microscope using anti-Flag (M2): F1804 Sigma-Aldrich, anti-calnexin C5C9 CST antibodies.

### EM particle measurement

Nanoparticle size measurements were performed using ImageJ.

### Dark field microscopy

Imaging was performed on a NikonA1 microscope with a 1.2-1.4NA oil darkfield condenser. Analysis was performed with ImageJ.

### Holotomography

Holotomography quantitative phase imaging and data analysis was used using a Tomocube system http://www.tomocube.com/.

### Cloning and expression

De novo *S. lessoniana* RNA transcriptomes were generated using PE250 Illumina sequencing and assembled with Trinity (13). Full length cDNAs cloned with Takara Cat.# 634858 and confirmed by Sanger sequencing and gene synthesized by IDT. Reflectin proteins were cloned into pBOB-CAG, transfected by lipofectamine, 2000, 3000 or LTX, and analyzed 36-72 hrs post transfection.

## Acknowledgements

JO, NUT & GMP were funded by a WM Keck foundation medical research grant and a Salk Institute Jacobs innovation grant. We would like to thank Tony Hunter for guidance and support. LH is at the Razavi Newman Integrative Genomics and Bioinformatics Core Facility of the Salk Institute with funding from NIH-NCI CCSG: P30 014195, and the Helmsley Trust. We would like to acknowledge the help and the expertise of the Salk Institute biophotonics core facility for assistance, in particular, Sarah Dunn, Sammy Weiser-Novak, Leo Andrade and Uri Manor, and Sadaaki Kayama for *S. lessoniana* field work.

## Author Contributions

GMP conceived the project. JO, GMP cloned reflectin genes. JO performed all the microscopy. NUT provided molecular biology technical assistance. RA and AM organized molecular biology logistics for the work in Japan. YI and SI organized field work. YI, GMP, SI, and JO did field work. JO, GMP and CG performed FACS experiments. GMP and JO designed FACS and microscopy experiments. AM, GMP and JO designed holotomography experiments. LH assembled the transcriptomes and identified reflectin genes. IMV gave molecular biology advice. AM gave microscopy advice. YI gave cephalopod biology advice. IMV and GMP obtained funding for the project. GMP and IMV wrote the manuscript.

P.S. During the preparation of this manuscript work from the Gorodetzky lab showed similar & complementary results using the *Doryteuthis opalescens* reflectin A1 protein. Chatterjee, *et al. Nat Commun* **11**,2708 (2020). https://doi.org/10.1038/s41467-020-16151-6

